# A novel single-cell based method for breast cancer prognosis

**DOI:** 10.1101/2020.04.26.062794

**Authors:** Xiaomei Li, Lin Liu, Greg Goodall, Andreas Schreiber, Taosheng Xu, Jiuyong Li, Thuc Duy Le

## Abstract

Breast cancer prognosis is challenging due to the heterogeneity of the disease. Various computational methods using bulk RNA-seq data have been proposed for breast cancer prognosis. However, these methods suffer from limited performances or ambiguous biological relevance, as a result of the neglect of intra-tumor heterogeneity. Recently, single cell RNA-sequencing (scRNA-seq) has emerged for studying tumor heterogeneity at cellular levels. In this paper, we propose a novel method, *scPrognosis*, to improve breast cancer prognosis with scRNA-seq data. *scPrognosis* uses the scRNA-seq data of the biological process Epithelial-to-Mesenchymal Transition (EMT). It firstly infers the EMT pseudotime and a dynamic gene co-expression network, then uses an integrative model to select genes important in EMT based on their expression variation and differentiation in different stages of EMT, and their roles in the dynamic gene co-expression network. To validate and apply the selected signatures to breast cancer prognosis, we use them as the features to build a prediction model with bulk RNA-seq data. The experimental results show that *scPrognosis* outperforms other benchmark breast cancer prognosis methods that use bulk RNA-seq data. Moreover, the dynamic changes in the expression of the selected signature genes in EMT may provide clues to the link between EMT and clinical outcomes of breast cancer. *scPrognosis* will also be useful when applied to scRNA-seq datasets of different biological processes other than EMT.

**Author summary:** Various computational methods have been developed for breast cancer prognosis. However, those methods mainly use the gene expression data generated by the bulk RNA sequencing techniques, which average the expression level of a gene across different cell types. As breast cancer is a heterogenous disease, the bulk gene expression may not be the ideal resource for cancer prognosis. In this study, we propose a novel method to improve breast cancer prognosis using scRNA-seq data. The proposed method has been applied to the EMT scRNA-seq dataset for identifying breast cancer signatures for prognosis. In comparison with existing bulk expression data based methods in breast cancer prognosis, our method shows a better performance. Our single-cell-based signatures provide clues to the relation between EMT and clinical outcomes of breast cancer. In addition, the proposed method can also be useful when applied to scRNA-seq datasets of different biological processes other than EMT.

## Introduction

Cancer prognosis plays an important role in clinical decision making. Traditionally, cancer prognosis is based on several clinical and pathological variables such as tumor size, lymph node status, histological grades, and so on [1]. However, these clinicopathological factors are insufficient for cancer prognosis because cancer is heterogeneous at the molecular (e.g., genes) level. Hence, recent clinical guidelines have highlighted the importance of using multi-gene tests to select patients who should receive adjuvant therapies [2]. The multiple genes in the tests are known as cancer signatures, which are crucial to cancer prognosis. Cancer signatures can be identified by in vivo biological experiments. For example, the *LM* method [3] analyzed transcriptomics in the cell lines and chose 54 genes associated with lung metastagenicity and virulence. However, these experiments cannot be done on human beings. Meanwhile, experiments on animals would not guarantee that the same conclusion can be drawn for humans. Therefore, computational methods are needed to identify cancer signatures from existing data, including gene expression data and clinical data.

Computational methods for breast cancer prognosis have shown some successes. Generally, these methods select the prognostic genes from a large number of human genes and then train survival models based on the selected genes. For instance, *PAM50* starts with an extended intrinsic gene set from previous studies, then selects genes based on their contributions in terms of distinguishing the five intrinsic breast cancer subtypes [4]. The *RS* method selects 16 cancer signatures from 250 published candidate genes [5]. *Mamma* [6] and *GGI97* [7] use a statistical test to choose the genes which differentially express between two distinct groups of tumors. Most of these methods use supervised algorithms to select the candidate genes and only *GGI97* ranks genes based on the similarities between gene expression profiles and tumor histologic grades. Based on the selected genes, most methods train linear regression models to predict the outcomes of the new coming patients. The clinical benefits of these prognostic genes for breast cancer are well studied on the traditional transcriptomics data, and some of the methods have approved by the Food and Drug Administration for commercial use [2].

The common feature of existing computational methods for breast cancer prognosis is that they are based on bulk RNA-seq data, which can lead to the following problems. Firstly, different tumor samples in bulk RNA-seq data have different proportions of cancer cells (named tumor purities) that can bias the results of these methods [8]. The traditional RNA sequencing technology measures the average expression levels of genes for an ensemble of cells from a tumor sample to obtain the so called bulk RNA-seq data. As a solid tumor tissue is a mixture of normal and cancer cells, the bulk RNA-seq data hence contain mixed signals and the non-cancerous components may have influences on genomic analysis of the bulk RNA-seq data or even bias the results [8]. There are works to uncover tumor purity and correct the bias in the detecting of differential genes [9] and identification of cancer subtypes [10]. It has been shown that differentially expressed genes and cancer subtypes are crucial to the selection of cancer signatures.

Secondly, with bulk RNA-seq data, we may not able to determine how gene signatures are related to cell level perturbation during cancer progression. Increasing evidence shows that the expression patterns of genes are heterogeneous from cell to cell [11]. These stochastic expression patterns trigger cell fate decisions and can affect cancer initiation and progress. However, based on the bulk RNA-seq data, the existing cancer prognosis methods cannot determine the correlation between clinical outcomes and dynamic gene behaviors along cellular trajectory.

Single cell RNA sequencing (scRNA-seq) has emerged recently and has many advantages over bulk RNA sequencing. Firstly, scRNA-seq does not have the tumor purity problem because it is possible to discover the existence of the micro-environment cell populations from scRNA-Seq data (See a review on this in [12]). Secondly, scRNA-seq is a powerful method to comprehensively characterize the cellular perturbation or stages within tissues [13] as it measures the expression of genes in individual cells. Additionally, scRNA-seq trajectory methods can provide a precise understanding of dynamic cell fate differentiation (See a systematic comparison in [14]). Through continuous cell stages along the pseudo-trajectory, we can observe the stochastic nature of gene expression [15]. Currently, scRNA-seq data are mostly used to detect cell types or to find novel biomarkers. As far as we know, there has been no work conducted on using scRNA-seq data to improve breast cancer prognosis.

In this work, we develop a novel method called *scPrognosis* to use scRNA-seq data to identify breast cancer signatures. Epithelial to Mesenchymal Transition (EMT) is a biological process associated with carcinogenesis, invasion, metastasis, and resistance to therapy in cancer [16]. We hypothesize that genes that play an important role in EMT are associated with breast cancer prognosis. Hence, we use an EMT scRNA-seq dataset for identifying the breast cancer signatures for prognosis. To fully exploit the scRNA-seq data towards optimal identification of breast cancer signatures, we propose to assess the importance of genes in the EMT process by integrating the following three measures: (1) their median absolute deviation in expression level; (2) their differentiation in different stages of EMT; (3) their roles in the dynamic gene co-expression network in EMT. *scPrognosis* uses a linear model to integrate the three measures for inferring breast cancer signatures. The significant difference between our method and the bulk RNA-seq data based methods is that we reconstruct the pseudotemporal trajectory known as pseudotime of cells [17] in EMT and incorporate this information into differential gene expression analysis and dynamic gene co-expression network construction. To validate the prognostic ability of these discovered gene signatures and apply them to breast cancer prognosis, we use them to build prediction models using bulk RNA-seq data as the data contains matched clinical information (and there are no single cell data with matched clinical information available). We apply *scPrognosis* to four independent bulk breast cancer datasets, ranging from about 200 to 1200 patients. The experimental results show that *scPrognosis* improves cancer prognosis compared with other benchmark breast cancer prognosis methods based on bulk RNA-seq data. A significant portion of the discovered prognostic genes is proved to be associated with breast cancer prognosis. Moreover, the dynamic changes in the expression trends of the genes provide clues to the link between EMT transition and clinical outcomes of breast cancer.

## Materials and methods

### Overview of *scPrognosis*

*scPrognosis* contains five steps as depicted in Fig 1. In step 1, *MAGIC* [18] and a gene filter are used to pre-process the noisy and high-dimensional scRNA-seq data. In step 2,EMT pseudotime, pseudotime series gene expression data, and dynamic gene co-expression network are inferred from the scRNA-seq data. In this step, firstly VIM gene expression level and pseudotemporal trajectory estimated by the *Wanderlust* algorithm [19] are used to identify EMT pseudotime for all cells in the scRNA-seq dataset. The EMT pseudotime describes the gradual transition of the single-cell transcriptome during the EMT transition process and helps to study gene expression dynamics in different EMT transition stages. Secondly, pseudotime series gene expression data is obtained by ordering cells in the scRNA-seq dataset from epithelial stage to mesenchymal stage according to the EMT pseudotime. Thirdly, from the ordered scRNA-seq data, a dynamic gene co-expression network is constructed by using the *LEAP* R package [20]. In step 3, based on the ordered scRNA-seq data, three methods are adopted to obtain the different gene ranking measures, including Median Absolute deviation (*MAD*), *switchde* [15] and Google *PageRank*. *MAD* and *switchde* are used to compute gene importance based on their expression level. Google *PageRank* ranks genes based on their roles in the dynamic gene co-expression network. In step 4, we integrate the three different rankings obtained in step 3 to prioritize genes. In step 5, the top N ranked genes are selected as signatures to predict the survival outcomes of breast cancer patients in bulk RNA-seq data. Details of each step are described in the following sub-sections.

**Fig 1.**
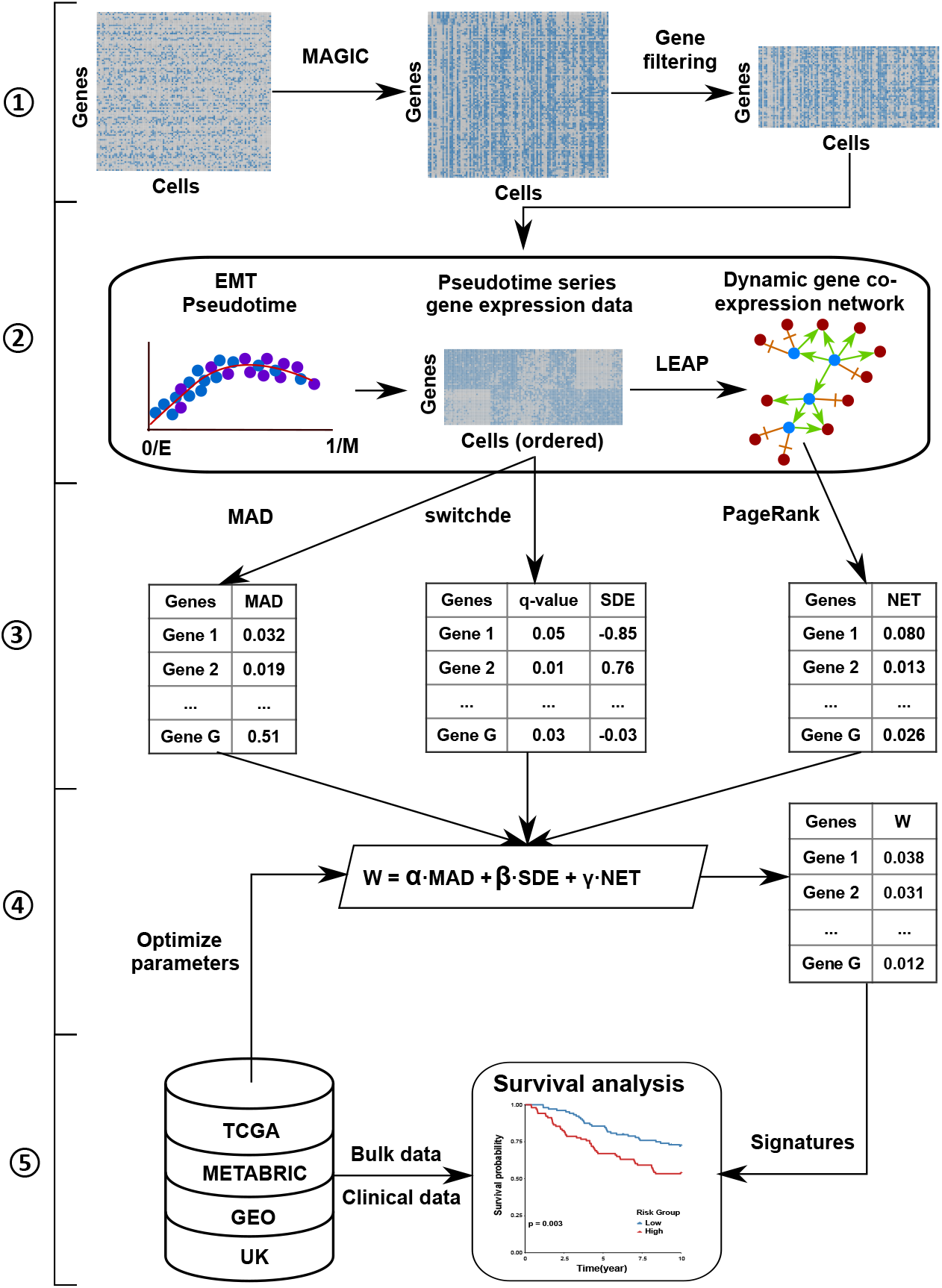
Workflow of the proposed *scPrognosis* framework. There are five main steps in *scPrognosis*, including: ➀ Pre-processing scRNA-seq data; ➁ Inferring EMT pseudotime, pseudotime series gene expression data, and dynamic gene co-expression network from the filtered scRNA-seq data; ➂ Ranking genes by three measurements; ➃ Prioritizing genes via an integrative model; ➄ Cancer prognosis using the top N ranked genes. The first four steps are based on scRNA-seq data while the last step uses bulk RNA-seq data to select parameters.

### Pre-processing scRNA-seq data

In the first step, *scPrognosis* pre-processes the input scRNA-seq dataset. The scRNA-seq dataset is a data matrix with *G* rows and *C* columns, where each column stores the expression levels of *G* genes in a single cell. Due to the low amounts of transcripts in a cell, an expressed gene may not be detected during sequencing with current scRNA-seq technology. This can lead to missing values of expressed genes, which is called the “dropout” phenomenon. For example, scRNA-seq data by the *inDrops* platform only have about 30% effective reads for each cell. The dropout events can lead to significant bias in gene-gene relationships and other downstream analyses [18].

*MAGIC* [18] is a method to denoise scRNA-seq data and impute the missing gene expression profiles. To overcome the sparsity and noise of the raw count matrix, *MAGIC* uses PCA (principal component analysis) components to calculate cell-cell distance matrix. The distance matrix is converted to a cell-cell affinity (similarity) matrix by an adaptive Gaussian kernel method. The affinity matrix is symmetrized and Markov-normalized to construct a Markov transition matrix. The final denoised and imputed data matrix is obtained by multiplying the exponentiated Markov transition matrix by the raw count matrix. Based on the information sharing across similar cells, *MAGIC* recovers gene expression from the dropout and other sources of noise.

After the imputation, we filter out genes with low coverage rates and low expression levels because these genes are most likely not expressed. It is suggested that these genes should be removed when searching for discriminative genes in microarray data [21] and implementing the *switchde* method. More experimental details of *MAGIC* and the gene filter method are provided in Section 2 in S1 File.

### Inferring EMT pseudotime, pseudotime series gene expression data, and dynamic gene co-expression network

Recently, it has been proposed that EMT transition occurs through continuum stages and there are several intermediate stages known as hybrid (partial) epithelial/mesenchymal (E/M) stages. Interestingly, these hybrid E/M stages are stable and can be the endpoint of a transition [16]. This means that cells may not go through the whole EMT transition and stop at a hybrid E/M stage. Switch-like genes that are up- or down-regulation along the EMT trajectory may induce cells to undergo a transition from one hybrid E/M stage to another hybrid E/M stage. Applying the proposition to the continuum stages of EMT transition, we could characterize the nature of switch-like genes and dynamic gene-gene relationships along the EMT trajectory. The strength of switch-like changes and the importance of genes in the dynamic gene co-expression network will be used to rank genes in our methods.

In this step, we will firstly infer the EMT pseudotime, and then based on the obtained pseudotime, we construct the pseudotime series gene expression dataset from the scRNA-seq dataset, which will be used in Step 3 to capture the switch-like changes along the pseudotime. At the same time, we also construct the dynamic gene co-expression network based on the pseudotime series gene expression dataset.

Even we do not have the true time-series data of individual cells undergoing EMT transition, we still can use scRNA-seq trajectory method to infer pseudotime from static scRNA-seq data. We assume that the EMT trajectory is a linear topology of ordered single cells, and cells represent the entire developmental process from E to M, i.e. each cell in the ordered sequence represents a different stage of the E to M transition. The trajectory then provides an indication of the timeline of the EMT transition, known as the EMT pseudotime. The pseudotime can be obtained using different approaches. One simple way to approximate the EMT pseudotime from a static scRNA-seq dataset is to order cells by their expression values of VIM [18], and we denote this pseudotime as VIM-time. Another way to infer the EMT pseudotime from a scRNA-seq dataset is by using the *Wanderlust* algorithm [19]. *Wanderlust* is a graph-based method to infer a linear tread to recapitulate cell trajectory. *Wanderlust* converts scRNA-seq data into a k-nearest neighbor graph (k-NNG). In k-NNG, each node is a cell, and each cell is connected to *k* cells that have similar expression profiles. Then *Wanderlust* generates several l-out-of-k-nearest neighbor graphs (l-k-NNGs) by randomly keeping *l* of k-nearest neighbors for each node in the k-NNG. For each l-k-NNG, *Wanderlust* identifies a trajectory score for each cell using a repetitive randomized shortest path algorithm. The final trajectory is computed by the average over all graph trajectories. We use the final trajectory as the EMT pseudotime named W-time. All the parameter assignments of *Wanderlust* can be found in Section 2 in S1 File.

After obtaining the EMT pseudotime, we have a trajectory score ranging from 0 to 1 for each cell which indicates its developmental stage of the E to M transition. Therefore, the scRNA-seq dataset (a data matrix) can be converted to a pseudotime series gene expression dataset by sorting cells (columns) based on the EMT pseudotime.

Then we construct a dynamic gene co-expression network from the above obtained pseudotime series expression dataset. Each node of the network represents a gene in the dataset. To capture the dynamic regulatory relationship between two genes, we use *LEAP* (lag-based Expression Association Pseudotime-series) [20] package to determine if there is an edge between two nodes. Given *C* cells ordered by the EMT pseudotime, the *MAC counter()* function in *LEAP* calculates the maximum absolute correlation (MAC) between the two nodes across all the time lags *l* ∈ {0,1,…, *C*/3} using the pseudotime series expression data. If the MAC between two nodes *g* and *t_g_* is tested to be statistically significant, an edge is added from *g* → *t_g_*.

### The three measures for ranking genes

*scPrognosis* combines three measures to rank genes, including Median Absolute Deviation (MAD) of gene expression profiles, the Switch-like Differentiation of genes in different stages of EMT (SDE), and the roles played by genes in the gene co-expression NETwork in EMT (NET). In this step, *scPrognosis* calculates the three measures individually before they are integrated into the next step. In the following, we describe the details of calculating each of the measures.

Let (*e*_1_, *e*_2_,…, *e_C_*) represents the expression profile of a gene *g* ∈ {1,…, *G*}, where *C* is the number of cells. The MAD of the gene can be computed as:

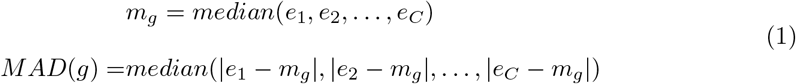

where *median()* is the function returning the median value of a given variable.

To calculate SDE, we use the software tool *switchde* [15] which can estimate the differentiation of switch-like genes in different stages of EMT. *switchde* defines a sigmoid function as shown in Eq 2 to fit the profile of a gene *g* with regard to a pseudotime *t_c_* (c is the index of a cell and *c* ∈ {1,…, *C*}). In Eq 2, 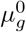, *k_g_* and 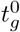 are the average peak expression value, the active strength and the active time of *g*. *k_g_* presents how quickly the gene *g* is up or down regulated along the pseudotime. We define *SDE(g)* as the switch-like differential expression level of the gene *g*, and *SDE*(*g*) = *k_g_*.

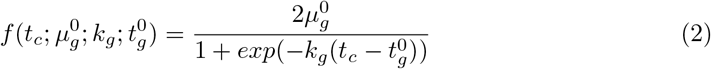

*switchde* adopts the gradient-based L-BFGS-B optimization algorithm [22] to obtain the maximum likelihood estimates of the parameters 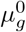, *k_g_*, and 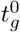. *switchde* also do the hypothesis testing associated with gene differential expression and adjust p-value by the Benjamini-Hochberg method.

To calculate NET, we follow the modified Google *PageRank* algorithm presented in [23]. The modified Google *PageRank* algorithm is used to calculate the regulatory importance of a gene in the dynamic gene co-expression network. Suppose there are *G* genes, the ranking of a gene *g* is defined as the following:

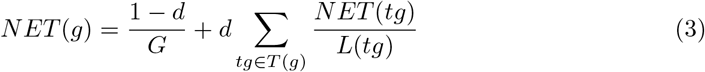

where *d* is the damping factor in *PageRank* and is set to 0.85 by default. *t_g_* is a target of *g* and we use *T(g)* to denote the set of all targets of *g*. *L(tg)* is the number of genes which regulate *t_g_*. From Eq 3, we can see that the rank of a gene depends on the rank of all its target genes. *NET(g)* is initialized to the same value for all *g*, and can be calculated using a iterative algorithm until it converges.

### Prioritizing genes via an integrative model

Although all the three measures are all associated with the clinical outcomes of cancer, none of the individual measure suffices to cancer prognosis. The expression variation (MAD) helps with distinguishing different cell populations. Genes with high expression variations are also of great clinical interest. The differentiation in different stages of EMT is corresponding to the gene behavior along the trajectory of EMT. SDE helps identify the genes that switch on and off alternatively during the trajectory to trigger EMT. The gene co-expression network is important for us to better understand the mechanisms of cell differentiation and carcinogenesis at a systems level. NET helps us discover hub regulatory genes that target the highest degree of a series of genes (called targets) in the network. It is believed that the hub regulatory genes are more closely related to cancer and have more biological significance compared with their targets [24]. Because each of them only reflects one aspect of the importance of a gene, and a combination of the three would be a more comprehensive measure. Therefore, we propose a linear model to integrate the three measures to obtain the final score for each gene.

Before integrating the three measures, we normalize them as follows:

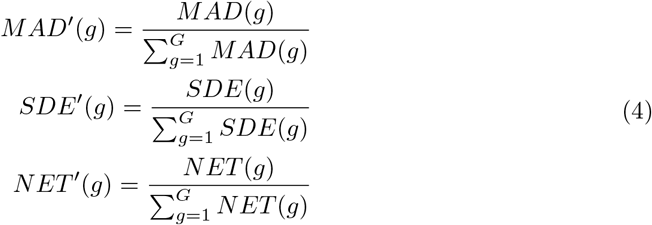

Then we integrate the normalised individual measures as follows.

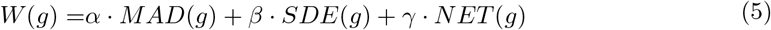

where *α*, *β*, and *γ* are the weights of MAD, SDE and NET respectively, *α* + *β* + *γ* = 1, and 0 ≤ *α, β, γ* ≤ 1. Then we rank the genes in descending order of the integrated measure. We use the grid search and cross validation methods to tune the weights of the linear model in the experiments. The optimal weights can lead to the best predictor of cancer prognosis on the bulk RNA-seq data.

### Cancer prognosis using the top N ranked genes

From the list of ranked genes obtained in Step 4, we select the top N ranked genes as cancer signatures. Then the Cox proportional hazards (PH) model [25] is trained based on these cancer signatures and bulk RNA-seq data. The PH model assumes that the effect of covariances on the survival outcomes is time-independent. Given survival time *t*, the general function of the PH model is defined as the following:

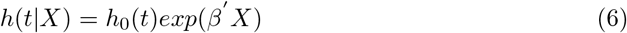

where *β*′ is a *N* × 1 vector that holds estimated regression coefficients, *X* is the expression data of the top N genes, and *h*_0_(*t*) is the baseline hazard function. The risk score of a new patient is calculated by

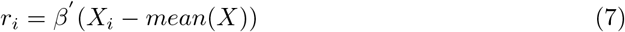

where *X_i_* is the expression data of the top N genes of the new patient *i*, and *mean()* is the function returning the average values of given data.

### Performance evaluation

#### Concordance index

(C-index) [26] is commonly used to validate the predictive ability of cancer prognostic models. Let *z_i_* and *r_i_* be the potential survival time and the risk score predicted using Eq 7 for patient *i*, respectively. C-index is equal to the concordance probability *P* (*r_i_* > *r_j_*|*z_i_* < *z_j_*) for a randomly selected pair of patients *i* and *j*. However, we cannot observe potential survival time for some patients who are lost to follow-up or event free at the end of a study (right censored). Hence the actually observed survival time *t_i_* = *min*(*z_i_*, *c_i_*), where *c_i_* is the potential right censoring time. Let *δ_i_* be the censoring status. An event (e.g. death or relapse) is developed within the study period when *δ_i_* = 1. For the right censoring data, C-index can be defined as the following:

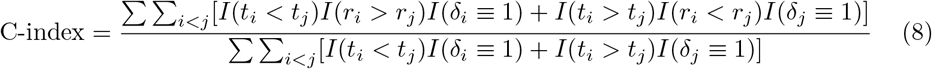

where *I()* is an indication function. C-index ranges from 0 to 1. The bigger the C-index is, the more accurate of a model will be.

#### Hazard ratio

To assist clinicians in tailoring treatment strategy, we often need to stratify patients into the high-risk group and the low-risk group via dichotomizing the predicted risk scores around their median value. Therefore, we need an accuracy measure to compare different methods. We use the hazard ratio (HR) as a accuracy measure, similar to other work [27]. We binarize the predicted risk scores to obtain the predicted groups *R* for patients. Then we estimate the risk difference between the two survival groups by Cox’s proportional hazards model as:

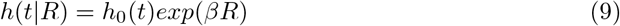

where *h*_0_(*t*) is the same as that in Eq 9. The quantity *exp*(*β*) is defined as HR, which indicates the risk difference between the two groups of patients. The larger the HR is, the larger discrimination between the low-and high-risk group becomes, and therefore the better the prediction method will be.

#### Kaplan-Meier survival curve

The Kaplan-Meier (KM) survival curve [28] combined with the Log-rank [29] test can identify whether the two risk groups show significantly different survival patterns. In the KM curve plot, the Y-axis is the probability of surviving in a given length of time, and the X-axis is survival time. The KM curves should have different characteristics and should not overlap for different groups predicted by a good method. The Log-rank test determines whether the survival curve estimated for each group is identical or not. If the p-value of the Log-rank rest is less than 0.05, the survival curves are statistically significantly different.

### Implementation

*scPrognosis* has been implemented using MATLAB and R packages. All the datasets and the R scripts to reproduce the results in this paper are available online at https://github.com/XiaomeiLi1/scPrognosis.

## Results

### Data sources and preparation

#### scRNA-seq data

In this paper, we use the scRNA-seq data of HMLE breast cancer cell lines from [18] to identify the EMT pseudotime for each cell and then select cancer signatures. The cells were stimulated with TGF-beta to induce EMT transition and the single-cell sequencing was performed using the *inDrops* platform. There are 28910 transcripts effectively measured in 7523 single cells. The scRNA-seq data can be download from the Gene Expression Omnibus (GEO) database (https://www.ncbi.nlm.nih.gov/geo/) under accession number GSE114397.

#### Bulk RNA-seq data

For training and validating the cancer prognosis model based on the selected signatures, we use bulk RNA-seq data of 2979 breast cancer patients from four different repositories, including TCGA (753 samples), METABRIC (1283 samples), GEO (736 samples) and UK (207 samples). Most of the breast cancer samples possess detailed clinical data, such as age, nodal, stage, grade, survival time, and event status. The TCGA and METABRIC datasets contain both overall survival time (OS) and relapse-free survival (RF) endpoints. The GEO and UK datasets only have the endpoints of relapse-free survival. The TCGA dataset was downloaded from the TCGA data portal (http://firebrowse.org/) and the dataset consists of level 3 mRNA expression data of primary breast cancer. The METABRIC dataset [30] was downloaded from the European Genome-phenome Archive (https://www.ebi.ac.uk/ega/ accession number EGAS00000000083, approval needed). The GEO dataset consists of 5 datasets: GSE12276 (204 samples), GSE19615 (115 samples), GSE20711 (88 samples), GSE21653 (252 samples) and GSE9195 (77 samples). We merge the five GEO datasets into a bigger dataset and adjusted the batch effects by the *ComBat* algorithm from the *sva* library [31]. The UK (known as GSE22219) dataset contains 207 early-invasive breast cancer cases with complete follow-up clinical data in 10 years. Both the GEO and UK datasets were downloaded from the Gene Expression Omnibus repository (https://www.ncbi.nlm.nih.gov/geo/). We summarize the details of these bulk RNA-seq datasets in Table 1.

**Table 1.** The description of bulk RNA-seq datasets.

| Dataset | Platform | Sample size | #transcripts |
| --- | --- | --- | --- |
| TCGA | Illumina RNA-seq | 753 | 13088 |
| METABRIC | Illumina RNA-seq | 1283 | 25191 |
| GEO | Affymetrix microarray | 736 | 18503 |
| UK | Illumina microarray | 207 | 22172 |

### *scPrognosis* is better than benchmark methods for risk score *prediction*

As discussed in the previous section, the SDE and NET measures have a considerable dependency on the pseudotime. We investigate the performance of two versions of *scPrognosis* based on different pseudotime, including VIM-time and W-time which are based on the expression profile of gene *VIM* and the *Wanderlust* algorithm, respectively. Section 2 in S1 File has more experiment details of calculating VIM-time and W-time. We denote the two versions of the implementations of *scPrognosis* as scP.V and scP.W, corresponding to the use of VIM-time and W-time, respectively.

To illustrate that scRNA-seq data can help to select prognostic signatures of breast cancer, we choose six widely used breast cancer prognosis benchmark methods that are based on the signatures selected from bulk RNA-seq data. More information about the benchmark methods can be found in Section 1 and Table 1 in S1 File. We compare the performance of the two versions of *scPrognosis* (scP.V and scP.W) with the benchmark methods on the datasets listed in Table 1. We report the results on TCGA and METABRIC according to the overall survival (OS) and relapse-free (RF) time. For the GEO and UK datasets, we report the results on the relapse-free time. Table 2 shows the C-indices and the mean ranking scores of all the methods compared. The C-index shown is the average of 100 times 10-fold cross-validation on a dataset. Based on the C-indices, mean ranking scores are calculated by Friedman’s test, which is a two-way analysis of variance by ranks for related samples. scP.W is better than other methods since it wins three times. Compared to the benchmark methods, scP.W outperforms all the methods for the prediction of the risk of RF time on the TCGA and UK datasets. Moreover, from the mean ranking results, we can see that *scPrognosis* overall outperforms the benchmark methods.

**Table 2.** Performance Comparison of cancer prognosis using benchmark methods and the proposed methods (scP.V and scP.W).

|  | PAM50 | Mamma | RS | GGI97 | Endo | LM | scP.V | scP.W |
| --- | --- | --- | --- | --- | --- | --- | --- | --- |
| TCGA(OS) | 0.63 | 0.55 | 0.58 | 0.51 | 0.54 | **0.63** | **0.63** | 0.61 |
| TCGA(RF) | 0.61 | 0.58 | 0.63 | 0.57 | 0.59 | 0.56 | 0.61 | **0.68** |
| METABRIC(OS) | 0.59 | 0.57 | 0.58 | 0.55 | **0.60** | 0.58 | 0.59 | **0.60** |
| METABRIC(RF) | 0.63 | 0.61 | 0.63 | 0.58 | **0.65** | 0.61 | 0.63 | 0.64 |
| GEO | 0.53 | 0.54 | **0.58** | 0.48 | 0.55 | 0.51 | 0.56 | 0.56 |
| UK | 0.60 | 0.62 | 0.63 | 0.61 | 0.63 | 0.64 | 0.67 | **0.70** |
| Mean rank | 4.50 | 2.92 | 5.33 | 1.33 | 5.17 | 3.67 | 6.08 | **7.00** |

The top-performing result is highlighted for each dataset. The result of C-index is the average of 100 times 10-fold cross-validation on each dataset.

To test whether a method performs significantly better than the other, we conduct the Wilcoxon signed-rank test based on the C-indices of *scPrognosis* and the benchmark methods. The result shows that both scP.V and scP.W perform significantly better than *Mamma* (the p-values are 0.017 and 0.018, respectively) and *GGI97* (the p-values are and 0.017, respectively). scP.V significantly outperforms *LM* (p-value = 0.03) while scP.W is superior to *RS* significantly (p-value = 0.046). Moreover, as previously shown (Table 2), according to the mean ranking scores, our methods still marginally improve the other two methods, *PAM50* and *Endo*.

In summary, we only use scRNA-seq data to measure the importance of genes, whereas the benchmark methods use signatures directly obtained from breast cancer clinical data and prior knowledge. Even so, the results have shown that both scP.V and scP.W achieve better or competitive performance compared with the benchmark methods. This indicates scRNA-seq data can improve the performance of breast cancer prognosis, and the signatures of EMT potentially are high quality predictors for breast cancer prognosis.

### *scPrognosis* is better than benchmark methods for risk group prediction

In this section, we evaluate *scPrognosis* using the Hazard Ratio (HR) criterion, in comparison with the six benchmark methods. For each method, we stratify patients into two groups using the risk scores calculated by the method. If a patient’s risk score bigger than the median value the patient is put into the high-risk group, otherwise the patient is put into the low-risk group. The HRs for all the methods are reported in Table 3. We observe that the two versions of *scPrognosis* (scP.V and scP.W) win once and twice, respectively, but *PAM50*, *RS*, and *Endo* each wines once this time. Based on the mean ranking results, we can conclude that overall *scPrognosis* outperforms the benchmark methods in stratifying patients into two risk groups.

**Table 3.** Comparison of the performances of risk group predictions using benchmark methods and the proposed methods (scP.V and scP.W).

|  | PAM50 | Mamma | RS | GGI97 | Endo | LM | scP.V | scP.W |
| --- | --- | --- | --- | --- | --- | --- | --- | --- |
| TCGA(OS) | 1.45 | 0.95 | 1.33 | 0.87 | 0.87 | 1.43 | **1.76** | 1.32 |
| TCGA(RF) | **1.61** | 1.01 | 1.26 | 0.97 | 1.17 | 1.09 | 1.30 | 1.45 |
| METABRIC(OS) | 4.90 | 4.67 | 5.99 | 2.51 | 4.80 | 3.60 | 4.71 | **6.07** |
| METABRIC(RF) | 5.17 | 3.87 | 5.29 | 2.47 | **7.16** | 3.95 | 5.93 | 5.45 |
| GEO | 1.20 | 1.52 | **2.50** | 0.91 | 1.85 | 0.86 | 1.37 | 1.51 |
| UK | 1.54 | 1.72 | 1.62 | 1.68 | 2.17 | 1.74 | 2.90 | **3.27** |
| Mean rank | 4.83 | 3.33 | 5.33 | 1.58 | 5.25 | 3.33 | 6.00 | **6.33** |

The top-performing result is highlighted for each dataset. The result of hazard ratio is the average of 100 times 10-fold cross-validation on each dataset.

Then we use the Wilcoxon signed-rank test to test the significance of the results on the HR criterion. Again, both scP.V and scP.W have perform significantly better than *Mamma* (the p-values are 0.047 and 0.031, respectively), *GGI97* (the p-values are 0.016 and 0.016, respectively) and *LM* (the p-values are 0.016 and 0.030, respectively).

### Evaluation using independent test

According to the results in Tables 2 and 3, among the two different implementations of *scPrognosis*, scP.W outperforms scP.V. So we choose scP.W as our final method to identify breast cancer signatures. For further evaluating the robust of scP.W in breast cancer prognosis, we conduct independent tests on three bulk RNA-seq datasets. Due to the small sizes of the GEO, and UK datasets, we don’t train scP.W based on these datasets. Fig 2 shows the independent test results on TCGA when training on METABRIC. Figs 2(A) and 2(C) show the comparison of scP.W and the benchmark methods. In these two figures, the Y-axis is the C-index, and the X-axis is the category of methods. Based on C-index, scP.W achieves the best results in predicting overall survival and relapse-free survival time. Figs 2(B) and 2(D) are the KM curves and the Log-rank test of risk group prediction using scP.W on the TCGA dataset. The results show that scP.W successfully stratifies patients into two risk groups of relapse and overall survival. The p-values by the Log-rank test are less than 0.05, which indicates that two risk groups have significantly different survival patterns, and the high-risk group has lower survival probability than that of the low-risk group. The TCGA dataset is the second-largest dataset in breast cancer and widely used in breast cancer research. We report the comparison results of our method based on the TCGA dataset and the current breast cancer prognostic methods in Table 3 in S1 File. The results also show that scP.W achieves the best results in cancer prognosis.

**Fig 2.**
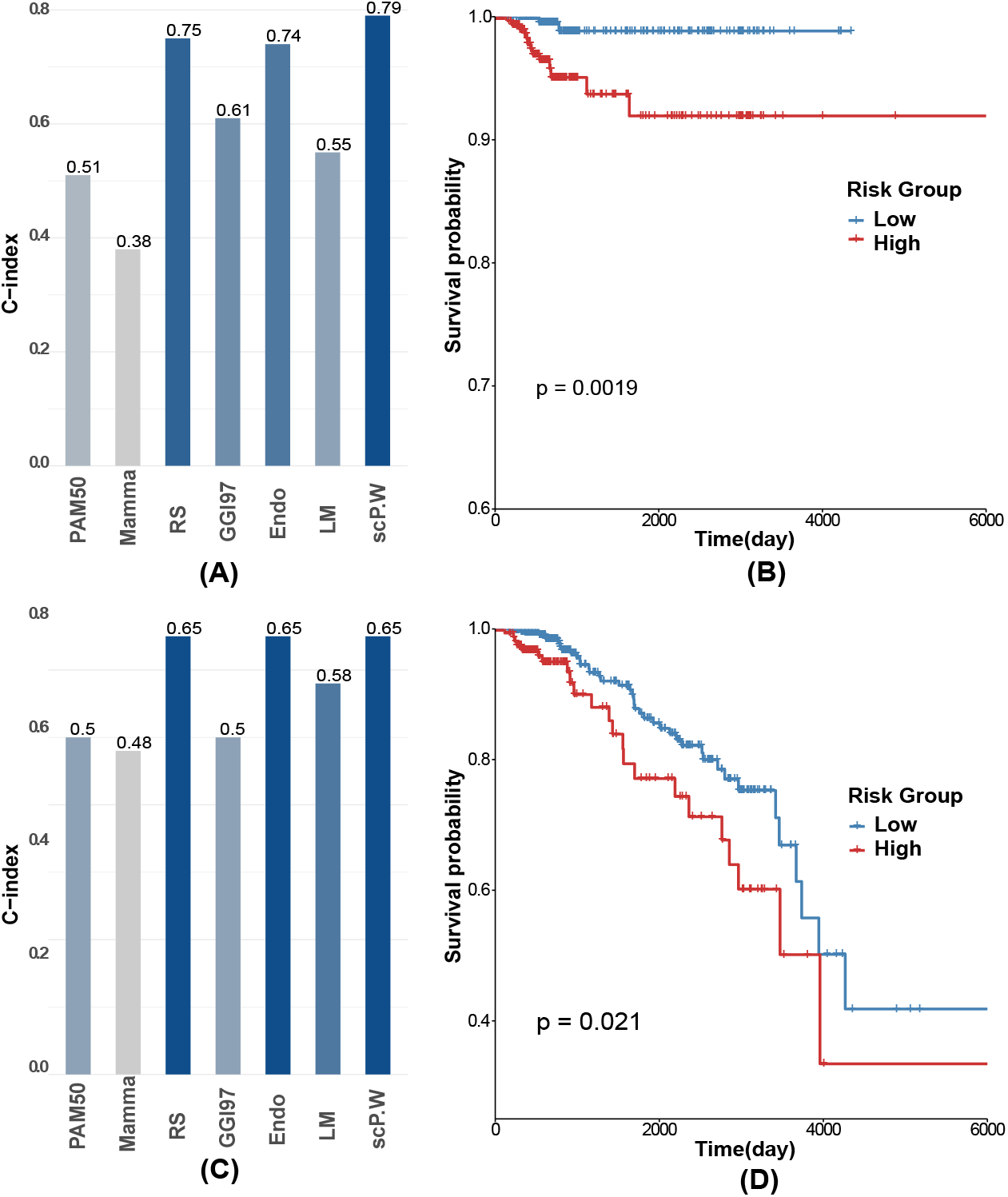
scP.W outperforms benchmark methods. (A) The bar chart of C-indices of scP.W and the benchmark methods on TCGA(OS); (B)The KM curve and Log-rank test of scP.W on TCGA(OS); (C)The bar chart of C-indices of scP.W and the benchmark methods on TCGA(RF); (D)The KM curve and Log-rank test of scP.W on TCGA(RF).

### Breast cancer signatures identified by scPrognosis

From the previous sections, we see the EMT signatures discovered by our methods are good breast cancer signatures too. To further validate these signatures, we compare the signatures discovered by our method with those discovered by benchmark methods. The EMT signatures are the top N ranked genes based on the scores calculated by Eq 5. Parameters N, *α*, *β*, and *γ* are determined by the 10-fold cross-validation results on bulk RNA-seq data.

scP.W selects 10 genes as breast cancer signatures, *KRT15*, *UBE2C*, *TOP2A*, *KRT6B*, *MKI67*, *HMGB2*, *ASPM*, *CDC20*, *KIF20A* and *CDK*, when trained on METABRIC. Comparing the 10 genes with the signatures used by the benchmark methods, we find 5 genes (*UBE2C*, *MKI67*, *ASPM*, *CDC20*, and *KIF20A*) showed up in one or more benchmark methods. *ASPM* is the common signature when scP.W is trained on TCGA and METABRIC. In our model, high *ASPM* levels are associated with adverse prognostic factors and shorter survival and relapse-free time. Recent evidence suggests that *ASPM* promotes prostate cancer stemness and progression and has important clinical and therapeutic significance [32]. Besides *ASPM*, other common signatures also have been proved to relate to breast cancer prognosis. For instance, high *UBE2C* expression is associated with poor prognosis in breast cancer, especially basal-like breast cancer [33]. *CDC20* over-expression means short-term breast cancer survival [34]. Fig 3 shows the diagram of overlapping genes among different methods. The diagram shows that a significant portion of the prognostic genes discovered by our method is overlapped with the current signatures of breast cancer prognosis. Though the clinical significance of the other five signature genes discovered by our method (*KRT15*, *TOP2A*, *KRT6B*, *HMGB2*, and *CDK1*) is not clear at present, they can be novel signatures for human breast cancer. There have been researches investigating the relationship between these genes and breast cancer. For example, *KRT6B* and *KRT15* were found to be the makers of basal-like breast cancers [35], and *TOP2A* expression levels were reported to have a significant association with metastasis-free survival in node-negative breast cancer [36].

**Fig 3.**
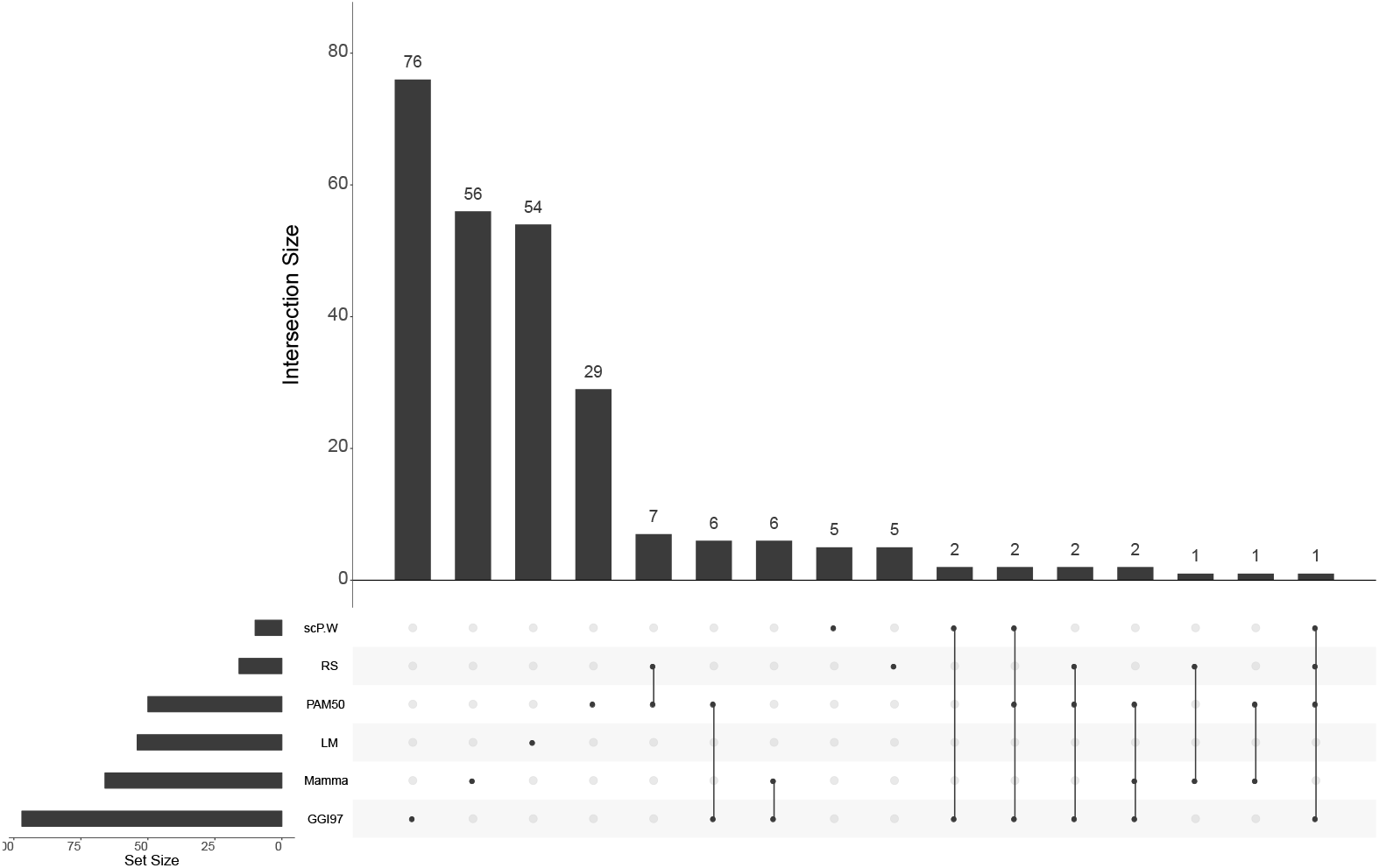
Overlap of signatures among different methods. The bottom left bar shows the number of signatures in each method. The dotted lines and the diagram on top show that the interaction overlaps among different methods. There are three genes (*UBE2C*, *MKI67*, and *CDC20*) in common with the scP.W, *PAM50*, and *GGI97*. Besides, scP.W has another two genes that only overlap with *GGI97* (*ASPM* and *KIF20A*). 5 out of 10 genes using scP.W have proved to associate with breast cancer prognosis.

### Correlation between EMT markers, breast cancer signatures and the EMT pseudotime

This paper is the first work to identify switch-like differential expression genes along the EMT pseudotime to understand their efficacy in deciphering the survival of breast cancer patients. No matter we use VIM-time or W-time, the models built have a good agreement on the performance. This is because W-time is highly related to the expression profile of VIM (the Pearson correlation is 0.46). For visualizing the dynamic behavior of genes in different stages of EMT, we divide W-time into 10 equal sized bins that present pseudo-stages of EMT. The expression level of a gene in a bin is calculated by the average of the profile during the time interval. Fig 4 shows the tendencies of the 10 cancer signatures along the pseudo-stages. We also plot the expression profiles of genes along W-time in Fig 1 in S1 File. Only *KRT15* and *KRT6B* are down-regulated by the EMT transition while other signatures do not vary at the E and M stages, but peak at the hybrid E/M stages. Recent experimental and theoretical evidence suggests that the hybrid E/M stages are stable phenotypes and is associated with aggressive tumor progression [37]. Our method demonstrates the relevance of the hybrid E/M phenotypes to patient survival in breast cancer.

**Fig 4.**
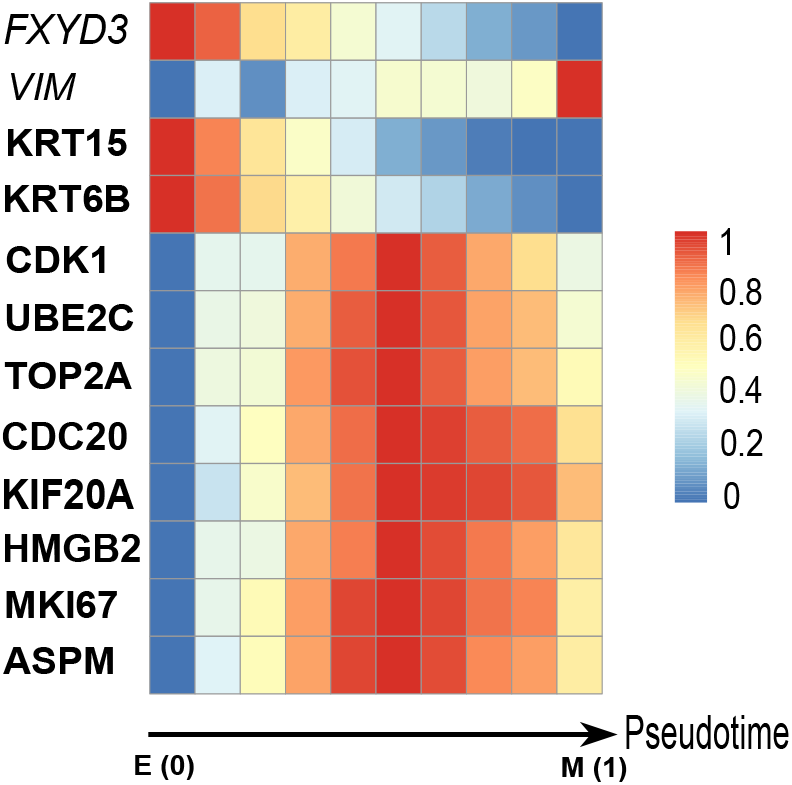
The heatmap of EMT markers and signatures along the EMT pseudotime. The bottom arrowed line is the W-time, and it indicates E stage, hybrid stages, and M stage from left to right on the line. The scale of the color bar on heatmap is from 0 to 1, and the color gradually changes from blue to red. The color of each cell of the colormap is based on the average expression level of a gene in a bin.

We also visualize the EMT markers’ dynamic behavior to determine whether the W-time could successfully model the cell evolved from the E stage to the M stage. The results from Fig 4 and Fig 1 in S1 File both show that the marker of epithelial (*FXYD3*) is down-regulated along W-time, while the marker of mesenchymal (*VIM*) is up-regulated along W-time. The tendencies of EMT markers along the W-time are consistent with prior knowledge that the expression of *VIM* increases while the expression of *FXYD3* decreases during the E to M transition. Therefore, W-time can successfully model the continuum of the E to M transition, and the results about the correlation between breast cancer signatures and EMT are reliable.

### Enrichment analysis of the signatures discovered by *scPrognosis*

We validate discovered breast cancer signature genes against the literature knowledge of pathways using the WikiPathways (http://www.wikipathways.org) platform [38]. The results in Table 4 show that the 10 signatures are highly relevant to the regulation of cancer. For instance, pathways 1, 2, 3, and 8 are direct pathways of cancer, and others are important pathways involved in the process of tumorigeneses.

**Table 4.** WikiPathways mapped pathways for the 10 breast cancer signatures.

| Id | Maps | P-value |
| --- | --- | --- |
| 1 | Gastric Cancer Network 1 WP2361 | 9.07E-05 |
| 2 | Gastric Cancer Network 2 WP2363 | 1.04E-04 |
| 3 | Retinoblastoma Gene in Cancer WP2446 | 9.33E-06 |
| 4 | Regulation of sister chromatid separation at the metaphase-anaphase transition WP4240 | 7.48E-03 |
| 5 | PPAR Alpha Pathway WP2878 | 1.29E-02 |
| 6 | Cell Cycle WP179 | 1.56E-03 |
| 7 | ATM Signaling Pathway WP2516 | 1.98E-02 |
| 8 | Integrated Cancer Pathway WP1971 | 2.18E-02 |
| 9 | ATM Signaling Network in Development and Disease WP3878 | 2.23E-02 |
| 10 | Regulation of Microtubule Cytoskeleton WP2038 | 2.28E-02 |

The pathways are highly relevant to the regulation of cancer.

We also conduct gene ontology enrichment analysis for the 10 breast cancer signatures. From Table 4 in S1 File, we can see that they are regulators of cell cycle progress and ubiquitin-protein ligase activities. Table 1 in S1 File shows that current signatures based on bulk RNA-seq data are also enriched in cell cycle regulation. Recent studies reveal the important roles of ubiquitin-protein ligase activity played in breast cancer [39, 40].

## Discussion and conclusion

Breast cancer is a complex disease caused by intricate genetic and molecular alterations. Thus traditional clinicopathological factors are not sufficient for the accurate prognosis of breast cancer. Recently, a wide range of computational methods have been proposed to identify multi-genes for breast cancer prognosis, and some of the methods have been approved for commercial use, including *PAM50*, *Mamma*, and *RS* test. These methods lead to a revolution in the breast cancer treatment paradigm. However, all of the progress in cancer prognosis has not been enough to overcome therapy resistance in breast cancer under current cancer therapeutics. Some tumor cells acquire resistance to targeted cancer therapy, which leads to worse survival of cancer patients. scRNA-seq can reveal genes that affect cell fate decision by monitoring the expression of genes in different cell states and sub-populations. In this paper, we use scRNA-seq data to detect signatures related to EMT that affect the clinical outcomes of breast cancer patients.

For almost two decades, the prospect that EMT may play an important role in tumor stemness, metastasis, and drug resistance has been vigorously debated. However, evidence demonstrating the prognosis power of EMT markers in breast cancer clinical studies has not been identified. Recently scRNA-seq is used to identify the continuum of EMT transition. We try to use the EMT scRNA-seq data to link the EMT related genes to breast cancer survival. To investigate how genes are related to cell level perturbation during EMT, we use the computational method *Wanderlust* to infer the EMT pseudotime. We integrate multiple measurements, MAD, SDE, and NET to measure the importance of a gene based on its expression variance, its dynamic differentiation, and its role in the dynamic gene co-expression network. We apply our method to four breast cancer cohorts. The experimental results illustrate that *scPrognosis* is more efficient than the benchmark methods based on bulk RNA-seq data and single-cell based methods only using individual measurements (Table 2 in S1 File). Our work also emphasizes the benefit of EMT mechanisms that incorporate background knowledge for identifying biologically relevant signatures of cancer prognosis. And the results show the good performance of the signatures in breast cancer prognosis.

Moreover, the results of *scPrognosis* may give us some clues for interpreting the EMT process. We look at the dynamic change of the gene expression along the EMT pseudotime. Interestingly, only two identified breast cancer signature genes are down-regulated along the EMT pseudotime, while the remaining genes peak at the intermediate of the E to M transition. These genes could be novel biomarkers for the hybrid E/M stages. We assume that the hybrid E/M stage is more relevant to patient survival as supported by the recent study in [41].

To identify the activity of EMT-related breast cancer signatures, we conduct a pathway analysis of the discovered breast cancer signatures. The results show that a significant number of the identified signatures are enriched in the pathways associated with cancer. Through the GO enrichment analysis, the signatures found by our method are closely related to the biological functions of cell cycle activity and ubiquitin-protein ligase activity, and the latter activity is not showing up in most of the current signatures.

However, there is no universal method that outperforms all the other methods. We still need to discover novel mechanisms involved in breast cancer progress, metastasis or resistance. In the future, our method can be extended to improve breast cancer prognosis by immune cell trajectories. Understanding immune cell development and response to disease is a crucial step for conquering cancer metastasis by immunotherapy. Recently there are some single cell experiments for investigating cellular dynamics in the context of immunology [42].

In conclusion, we have proposed a novel method *scPrognosis* for breast cancer prognosis based on scRNA-seq data. *scPrognosis* uses an integrative model to infer breast cancer signatures based on MAD, SDE, and NET measurements. We empirically compared our method with the existing methods on four breast cancer datasets. The results show that the scRNA-seq based method is a good and useful method for breast cancer prognosis. The signatures detected by our method show the link between EMT and the clinical outcomes of breast cancer, which may give some clues for current cancer therapeutics.

## Supporting information

**S1 File. Supplementary information**.

## Notes

### Competing Interest Statement

The authors have declared no competing interest.

## References

1. Goldhirsch A, Glick JH, Gelber RD, Coates AS, Thürlimann B, Senn HJ. Meeting highlights: international expert consensus on the primary therapy of early breast cancer 2005. Annals of oncology. 2005;16(10):1569–1583.

2. Duffy M, Harbeck N, Nap M, Molina R, Nicolini A, Senkus E, et al. Clinical use of biomarkers in breast cancer: Updated guidelines from the European Group on Tumor Markers (EGTM). European journal of cancer. 2017;75:284–298.

3. Minn AJ, Gupta GP, Siegel PM, Bos PD, Shu W, Giri DD, et al. Genes that mediate breast cancer metastasis to lung. Nature. 2005;436(7050):518.

4. Parker JS, Mullins M, Cheang MC, Leung S, Voduc D, Vickery T, et al. Supervised risk predictor of breast cancer based on intrinsic subtypes. Journal of clinical oncology. 2009;27(8):1160.

5. Paik S, Shak S, Tang G, Kim C, Baker J, Cronin M, et al. A multigene assay to predict recurrence of tamoxifen-treated, node-negative breast cancer. New England Journal of Medicine. 2004;351(27):2817–2826.

6. Van’t Veer LJ, Dai H, Van De Vijver MJ, He YD, Hart AA, Mao M, et al. Gene expression profiling predicts clinical outcome of breast cancer. nature. 2002;415(6871):530.

7. Sotiriou C, Wirapati P, Loi S, Harris A, Fox S, Smeds J, et al. Gene expression profiling in breast cancer: understanding the molecular basis of histologic grade to improve prognosis. Journal of the National Cancer Institute. 2006;98(4):262–272.

8. Aran D, Sirota M, Butte AJ. Systematic pan-cancer analysis of tumour purity. Nature communications. 2015;6:8971.

9. Yang J, Zhang W, Long H. DECtp: Calling differential gene expression between cancer and normal samples by integrating tumor purity information. Frontiers in genetics. 2018;9:321.

10. Zhang W, Feng H, Wu H, Zheng X. Accounting for tumor purity improves cancer subtype classification from DNA methylation data. Bioinformatics. 2017;33(17):2651–2657.

11. Huang S. Non-genetic heterogeneity of cells in development: more than just noise. Development. 2009;136(23):3853–3862.

12. Qi R, Ma A, Ma Q, Zou Q. Clustering and classification methods for single-cell RNA-sequencing data. Briefings in bioinformatics. 2019.

13. Tirosh I, Suvà ML. Deciphering Human Tumor Biology by Single-Cell Expression Profiling. Annual Review of Cancer Biology. 2019;3:151–166.

14. Saelens W, Cannoodt R, Todorov H, Saeys Y. A comparison of single-cell trajectory inference methods. Nature biotechnology. 2019;37(5):547.

15. Campbell KR, Yau C. switchde: inference of switch-like differential expression along single-cell trajectories. Bioinformatics. 2016;33(8):1241–1242.

16. Pastushenko I, Brisebarre A, Sifrim A, Fioramonti M, Revenco T, Boumahdi S, et al. Identification of the tumour transition states occurring during EMT. Nature. 2018;556(7702):463.

17. Trapnell C, Cacchiarelli D, Grimsby J, Pokharel P, Li S, Morse M, et al. Pseudo-temporal ordering of individual cells reveals dynamics and regulators of cell fate decisions. Nature biotechnology. 2014;32(4):381.

18. Van Dijk D, Sharma R, Nainys J, Yim K, Kathail P, Carr AJ, et al. Recovering gene interactions from single-cell data using data diffusion. Cell. 2018;174(3):716–729.

19. Bendall SC, Davis KL, Amir EaD, Tadmor MD, Simonds EF, Chen TJ, et al. Single-cell trajectory detection uncovers progression and regulatory coordination in human B cell development. Cell. 2014;157(3):714–725.

20. Specht AT, Li J. LEAP: constructing gene co-expression networks for single-cell RNA-sequencing data using pseudotime ordering. Bioinformatics. 2016;33(5):764–766.

21. Winter C, Kristiansen G, Kersting S, Roy J, Aust D, Knösel T, et al. Google goes cancer: improving outcome prediction for cancer patients by network-based ranking of marker genes. PLoS computational biology. 2012;8(5).

22. Byrd RH, Lu P, Nocedal J, Zhu C. A limited memory algorithm for bound constrained optimization. SIAM Journal on scientific computing. 1995;16(5):1190–1208.

23. Xu T, Le TD, Liu L, Wang R, Sun B, Li J. Identifying cancer subtypes from mirna-tf-mrna regulatory networks and expression data. PloS one. 2016;11(4):e0152792.

24. Chen J, Yu L, Zhang S, Chen X. Network analysis-based approach for exploring the potential diagnostic biomarkers of acute myocardial infarction. Frontiers in physiology. 2016;7:615.

25. Cox DR. Regression models and life-tables. In: Breakthroughs in statistics. Springer; 1992. p. 527–541.

26. Harrell FE, Lee KL, Mark DB. Multivariable prognostic models: issues in developing models, evaluating assumptions and adequacy, and measuring and reducing errors. Statistics in medicine. 1996;15(4):361–387.

27. Haibe-Kains B, Desmedt C, Sotiriou C, Bontempi G. A comparative study of survival models for breast cancer prognostication based on microarray data: does a single gene beat them all? Bioinformatics. 2008;24(19):2200–2208.

28. Rich JT, Neely JG, Paniello RC, Voelker CC, Nussenbaum B, Wang EW. A practical guide to understanding Kaplan-Meier curves. Otolaryngology—Head and Neck Surgery. 2010;143(3):331–336.

29. Bland JM, Altman DG. The logrank test. Bmj. 2004;328(7447):1073.

30. Curtis C, Shah SP, Chin SF, Turashvili G, Rueda OM, Dunning MJ, et al. The genomic and transcriptomic architecture of 2,000 breast tumours reveals novel subgroups. Nature. 2012;486(7403):346.

31. Leek JT, Johnson WE, Parker HS, Jaffe AE, Storey JD. The sva package for removing batch effects and other unwanted variation in high-throughput experiments. Bioinformatics. 2012;28(6):882–883.

32. Pai VC, Hsu CC, Chan TS, Liao WY, Chuu CP, Chen WY, et al. ASPM promotes prostate cancer stemness and progression by augmenting Wnt-Dvl-3-beta-catenin signaling. Oncogene. 2018;38:1340–53.

33. Qin T, Huang G, Chi L, Sui S, Song C, Li N, et al. Exceptionally high UBE2C expression is a unique phenomenon in basal-like type breast cancer and is regulated by BRCA1. Biomedicine & Pharmacotherapy. 2017;95:649–655.

34. Karra H, Repo H, Ahonen I, Löyttyniemi E, Pitkänen R, Lintunen M, et al. Cdc20 and securin overexpression predict short-term breast cancer survival. British journal of cancer. 2014;110(12):2905.

35. Charafe-Jauffret E, Ginestier C, Monville F, Finetti P, Adeläıde J, Cervera N, et al. Gene expression profiling of breast cell lines identifies potential new basal markers. Oncogene. 2006;25(15):2273.

36. Brase JC, Schmidt M, Fischbach T, Sültmann H, Bojar H, Koelbl H, et al. ERBB2 and TOP2A in breast cancer: a comprehensive analysis of gene amplification, RNA levels, and protein expression and their influence on prognosis and prediction. Clinical Cancer Research. 2010;16(8):2391–2401.

37. Kröger C, Afeyan A, Mraz J, Eaton EN, Reinhardt F, Khodor YL, et al. Acquisition of a hybrid E/M state is essential for tumorigenicity of basal breast cancer cells. Proceedings of the National Academy of Sciences. 2019;116(15):7353–7362.

38. Kutmon M, Riutta A, Nunes N, Hanspers K, Willighagen EL, Bohler A, et al. WikiPathways: capturing the full diversity of pathway knowledge. Nucleic acids research. 2015;44(D1):D488–D494.

39. Liao L, Song M, Li X, Tang L, Zhang T, Zhang L, et al. E3 ubiquitin ligase UBR5 drives the growth and metastasis of triple-negative breast cancer. Cancer research. 2017;77(8):2090–2101.

40. Ka WH, Cho SK, Chun BN, Byun SY, Ahn JC. The ubiquitin ligase COP1 regulates cell cycle and apoptosis by affecting p53 function in human breast cancer cell lines. Breast Cancer. 2018;25(5):529–538.

41. George JT, Jolly MK, Xu S, Somarelli JA, Levine H. Survival outcomes in cancer patients predicted by a partial EMT gene expression scoring metric. Cancer research. 2017;77(22):6415–6428.

42. Kunz DJ, Gomes T, James KR. Immune cell dynamics unfolded by single-cell technologies. Frontiers in immunology. 2018;9:1435.

